# Trajectories of brain organisation transition from predicting externalising to internalising symptoms across adolescence

**DOI:** 10.64898/2026.05.18.724880

**Authors:** Antoine Bernas, Linda Schlüter, Tobias Banaschewski, Arun L.W. Bokde, Rüdiger Brühl, Sylvane Desrivières, Herta Flor, Hugh Garavan, Penny Gowland, Antoine Grigis, Andreas Heinz, Herve Lemaitre, Jean-Luc Martinot, Marie-Laure Paillère Martinot, Eric Artigues, Frauke Nees, Dimitri Papadopoulos Orfanos, Tomáš Paus, Luise Poustka, Michael N. Smolka, Nathalie Holz, Nilakshi Vaidya, Henrik Walter, Robert Whelan, Paul Wirsching, Gunter Schumann, Andre Marquand

## Abstract

Understanding the dynamics of brain–behaviour relationships during adolescence is critical for elucidating the neurodevelopmental basis of mental health. Leveraging two large-scale longitudinal cohorts—the Adolescent Brain Cognitive Development (ABCD) and IMAGEN studies, comprising over 10,000 participants aged 10 to 22 years with six waves of multimodal neuroimaging and behavioural data, we applied multi-view sparse canonical correlation analysis to investigate evolving associations between structural MRI, resting-state functional connectivity, and multi-domain behavioural measures. Our findings reveal four fundamental patterns of developmental reorganisation in brain-psychopathology relationships.

First, symptom profiles evolved from predominantly externalising features (aggression, attention problems) at ages 10-12 toward global psychopathology by age 14, then transitioned toward internalising features (e.g., anxiety, depression) by ages 19-22, reflecting fundamental shifts in vulnerability from behavioural dysregulation to affective disturbance.

Second, cortical thickness exhibited negative associations with externalising symptom profiles throughout development. During early adolescence (ages 10-14) this was driven by broadly distributed decreases across sensorimotor, temporal, visual, and cingulate regions alongside overall mean cortical thickness. After 14, this diffuse pattern shifted towards late maturing association cortices, notably the dorsolateral prefrontal and lateral temporal cortices.

Third, this was accompanied by subcortical effects that exhibited greater age-specificity: whilst cerebellar volume contributions were evident at most timepoints, basal ganglia volume influence was principally evident in early development (ages 10-12), with thalamic structures and global subcortical grey matter volume becoming dominant at age 14, marking a transition in which subcortical structures mediate psychopathology associations.

Fourth, functional connectivity showed a more dynamic developmental trajectory. During early adolescence, symptom associations were driven by positive connectivity between cognitive control and sensorimotor networks, whereas late adolescence exhibited predominantly positive connectivity patterns, transitioning from dense sensorimotor-frontoparietal configurations to more specific patterns involving the central executive and default-mode networks.

These findings fundamentally challenge static biomarker models, demonstrating that adolescent psychopathology reflects developmentally contingent brain-behaviour relationships rather than static neural markers. Age 14 emerges as a critical inflection point marked by convergent thalamic reconfiguration, global subcortical grey matter dominance, and symptom profile transitions. This work provides an empirical foundation for precision mental health strategies tailored to specific developmental windows, with implications for reducing psychiatric burden in youth.

## INTRODUCTION

Adolescence is a critical period of neurodevelopment marked by profound structural and functional brain changes that underpin cognitive, emotional, and behavioural maturation. Understanding how variations in brain structure and function relate to behavioural traits during this formative stage is essential for identifying both normative developmental trajectories and early markers of psychopathology, which manifest as deviations from an expected trajectory, especially since adolescence is known to herald the onset of many forms of psychopathology^1,2^

Multimodal neuroimaging, particularly structural and functional magnetic resonance imaging (sMRI and fMRI), can provide valuable insights into the structural and functional organisation of the adolescent brain and therefore provides an ideal tool to probe these developmental patterns. When paired with comprehensive behavioural assessments, these data can reveal meaningful complex brain–behaviour relationships. Many neuroimaging studies have aimed to identify the key alterations to an assumed developmental trajectory that give rise to psychopathology, both from theoretical^3,4^ and clinical perspectives^5–9^. Unifying principles have, however, remained elusive because of a high degree of heterogeneity across individuals and because the multifaceted alterations are likely reflected in multiple aspects of brain structure and function. Moreover, the complexity and high dimensionality of such datasets pose significant analytical challenges, requiring advanced statistical methods capable of integrating information across modalities.

Canonical Correlation Analysis (CCA) is a multivariate technique well-suited for uncovering patterns of covariation between sets of variables. In the context of neuroimaging, CCA has enabled the identification of latent modes of population-level covariation between imaging-derived features and behavioural measures. Many recent studies in children have demonstrated the utility of CCA in linking brain connectivity profiles to cognitive and psychological dimensions^7,8,10–12^. However, it is unclear how such brain-behaviour associations change throughout development, and brain–behaviour association studies are often limited to only two sets or views of data which is suboptimal considering the multimodal nature of psychopathology. For example, researchers have attempted to identify associations between resting-state fMRI connectivity and psychopathological symptoms^12^, while others have applied CCA using multimodal neuroimaging components and clinical symptoms^13,14^. Few studies have integrated multiple data modalities via multi-view CCA approaches^15^. Another important limitation of these studies is the lack of generalisability in the resulting models. It was recently demonstrated that, when using a CCA-based multivariate approach, it is particularly difficult to obtain generalisable brain-based biomarkers of child psychiatric symptoms^12,16^.

These factors highlight the critical need for longitudinal strategies that can accurately capture inter-individual variability in developmental dynamics. A major obstacle is that raw brain measures are confounded by factors such as age, sex, and scanning site, which can distort true brain–behaviour relationships. Normative modeling^17^—conceptually similar to paediatric growth charts but applied to brain features^18^—provides standardised scores that accommodate variation related to scanning site and underlying developmental trajectories. This essentially involves transforming raw brain features into deviations from a common reference model, which provides the ability to bind heterogeneous cohorts to a single common reference model and account for underlying changes related to developmental trajectories which may be sex- and region specific. This ultimately results in more reliable and trustworthy brain–behaviour relationships and facilitates comparisons across studies^19^. A second major limitation is related to the logistic difficulties in acquiring data that span the entire developmental period. This is crucial because associations with symptoms may change over time, particularly in view of the profound social, psychological and biological changes that occur during adolescence^20^.

In this study, we aimed to chart multimodal brain–behaviour associations throughout adolescence. To achieve this, we developed a novel variant of multi-view sparse canonical correlation analysis (msCCA) that enables us to examine multivariate relationships between structural MRI, resting-state functional connectivity, and a broad spectrum of behavioural and psychological measures, whilst accommodating missing data. We applied this to track brain symptom associations across two large independent adolescent cohorts: the Adolescent Brain Cognitive Development (ABCD) study and the IMAGEN study datasets^21,22^. Crucially, these cohorts together span a long developmental timeframe (9-22 years) that overlaps with the timeframe that most psychopathology emerges^3^. We used normative modelling to bind theses samples together and fit separate brain-behaviour association models at each timepoint across key stages of adolescent development, thereby charting the manner in which both the expression of symptoms and their underlying brain systems change throughout development.

## RESULTS

An overview of our analytic pipeline is given in Figure 1. First, we applied harmonised processing and careful data quality control to the ABCD and IMAGEN samples (see methods). Each dataset included three waves of brain and behavioural assessments collected across the key developmental window from ages 10 to 22 years with an overlap at 14 years. We then derived deviation scores (z-scores) with respect to large scale normative models (NMs) for brain structural (cortical thickness and subcortical volumes) and functional (resting-state network connectivity) phenotypes^18,19^ (Fig. 1a). To quantify psychopathology in ABCD, we used item-level scores from the Kiddie Schedule for Affective Disorders and Schizophrenia (K-SADS)^23^ and Child Behavioural Checklist (CBCL)^24^, whereas in IMAGEN, we used the Development And Well-Being Assessment (DAWBA)^25^ and Strengths and Difficulties Questionnaire (SDQ)^26^ scores. These instruments provide a broad and approximately equivalent coverage of the range of psychopathology. Although direct cross-dataset score harmonisation is often desirable, it can introduce bias^27^ and is not a prerequisite here given that each cohort and timepoint is analysed independently, preserving within-dataset statistical integrity. Therefore, we elected not to harmonise instruments in this study. Rather, cross-dataset convergence is evaluated at the interpretive level by comparing effect directions, magnitudes, and the specific symptom dimensions implicated, effectively treating replication across instruments as a strength rather than a limitation. Using these image-derived and behavioural phenotypes, we investigated the multimodal (structural, functional, and behavioural) associations at each timepoint by means of multi-view sparse CCA (Fig. 1b,c), which we embedded within a stability selection and out of sample testing framework for reliable estimates of generalisability (see methods). The resulting canonical correlations, their significance, and the different brain and behavioural patterns (or profiles) that the adolescents revealed in this study, are reported in the next sections (Fig. 1d).

**Figure 1.**
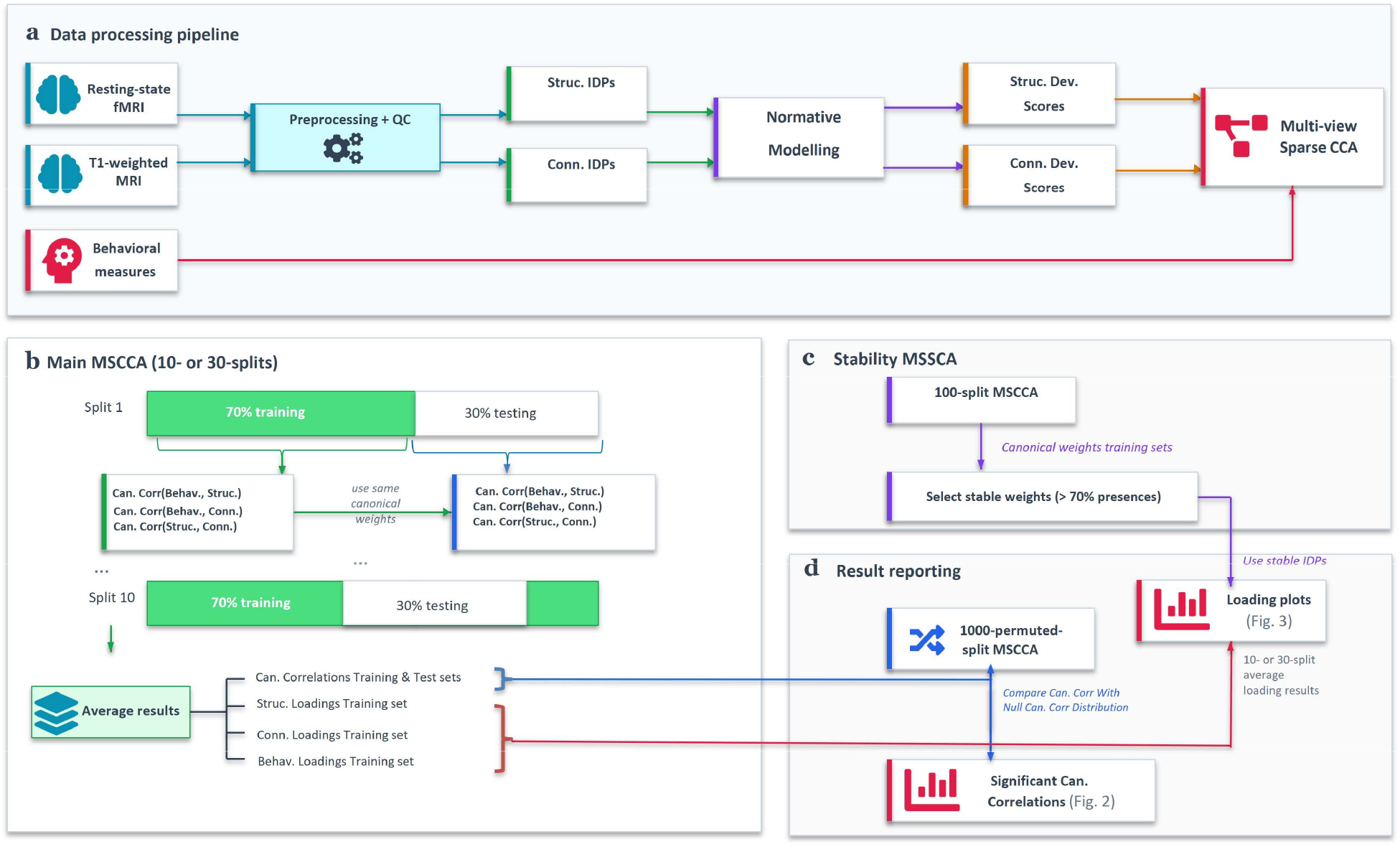
Analysis pipeline. **a**, Resting-state fMRI data and T1-weighted scans were preprocessed and quality controlled, which provided us with cortical thicknesses and subcortical volumes as structural (Struc.) and functional (Conn.). Structural and functional deviation (Dev.) z-scores were then predicted by fitting pre-trained normative models with these phenotypes. In parallel, the behavioural (Behav.; K-SADS, DAWBA bands, and SDQ) scores were normalised, and with the brain-derived z-scores underwent the multi-view sparse CCA (msCCA). **b**, Using the normalised data, we trained msCCA with 70% of the data in order to find canonical (Can.) weights, loadings, and Can. correlations (Corr.). Based on the training can. weights, we estimated the Can. Corr. of the test set (remaining 30% of the data). We repeated the approach 10 times and 30 times for ABCD and IMAGEN respectively (with randomised 70/30 train/test splits) and averaged the results (Can. Corr, and Loadings). **c**, Before statistically testing the canonical correlations and reporting the loadings, 100-split msCCA was performed (same way as the Main msCCA), and the IDP and Behav. that had non-null Can. weights, in at least 70 of the splits, were retained. **d**, We report, in this paper, the in-sample and out-of-sample significant Can. Corr. by use of permutation testing, and depict the stable (from the Stability msCCA) average loadings of the Main msCCA.

### Brain-symptom associations were evident at all timepoints across development

Across the different waves of the data, we observed significant brain-behaviour associations across all timepoints (Figure 2) that were also stable across 10 splits of the data and similar for training (70% in-sample) and test (30% out-of-sample) sets. To test the significance of the canonical correlations, we performed permutation testing (1000 permutations) and averaged the brain-behaviour correlations for symptoms with brain structure and symptoms with brain function. Note that we excluded the correlation between brain structure and function both from the optimisation and the assessment to prevent the model learning this trivial association. For more details, see Methods. In brief, we report significant associations at all timepoints; p-values are derived from the test set correlations. In ABCD, at 10 years old (ABCD_BL_), the canonical correlations in the training set (Rtr) = 0.12, test set (Rte) = 0.10 (p < 0.01). At 12 years (ABCD_FU1_), the associations were of similar magnitude Rtr/Rte = 0.13/0.09; p < 0.01. At age 14 (ABCD_FU2_), correlations were also significant: Rtr/Rte = 0.16/0.03; p < 0.01. Here, though, the weaker association in the test set (Rte = 0.03) suggests limited generalisability, likely attributable to the reduced sample size at this wave.

**Figure 2.**
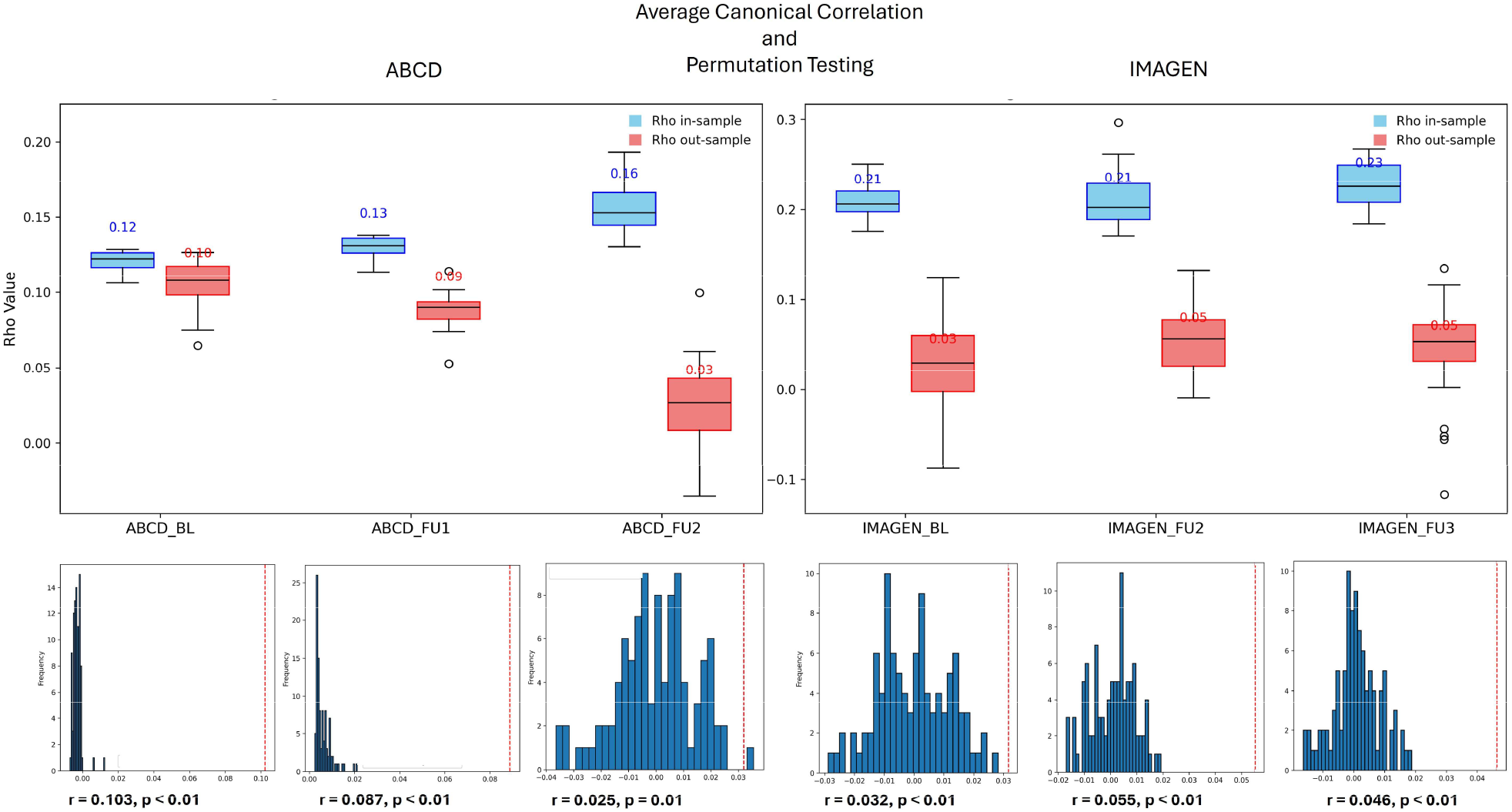
Significant canonical correlation boxplots. Average canonical correlation between the behavioural and brain-related canonical variates (top); and canonical correlation null distribution (bottom); for ABCD_BL_, ABCD_FU1_, and ABCD_FU2_, (left); and IMAGEN_BL_, IMAGEN_FU2_, and IMAGEN_FU3_ (right); In-sample (training) results in blue and Out-of-sample (test) results in red. The numbers displayed on the boxes are the 10-split (for ABCD) and 30-split (for IMAGEN) average correlations, while the boxes show the usual 25^th^, median, and 75^th^ centiles. Hollow circles = outliers, when correlation is outside of the whiskers (interquartile range x 1.5). r = out-of-sample average canonical correlation value (vertical red-dashed line); p = permutation test p-value.

We found largely similar associations across the IMAGEN dataset. Namely, significant associations between the brain deviation scores and symptoms at 14 years old (IMAGEN_BL_),: Rtr/Rte = 0.21/0.03 (p < 0.01), 19 years (IMAGEN_FU2_), Rtr/Rte = 0.20/0.05; p < 0.01 and 22 years (IMAGEN_FU3_), with Rtr/Rte = 0.19/0.05 (p < 0.01).

In summary, our msCCA models captured relevant patterns of brain–symptom association that remained statistically significant in the unseen validation/test splits of the data across all timepoints and across both datasets. Next, we examined the brain regions and symptoms underlying these associations. The brain structures and connectivity profiles driving these associations evolved across timepoints and datasets, as well as the associated symptom profiles, reflecting underlying developmental effects. These varied across different stages of adolescence, as described in the following sections and in Figure 3. It is worth noting that we present only the first canonical mode in the main text. However, all of the second- and some of the third modes also showed statistically significant canonical correlations; see supplementary figure S1.

**Figure 3.**
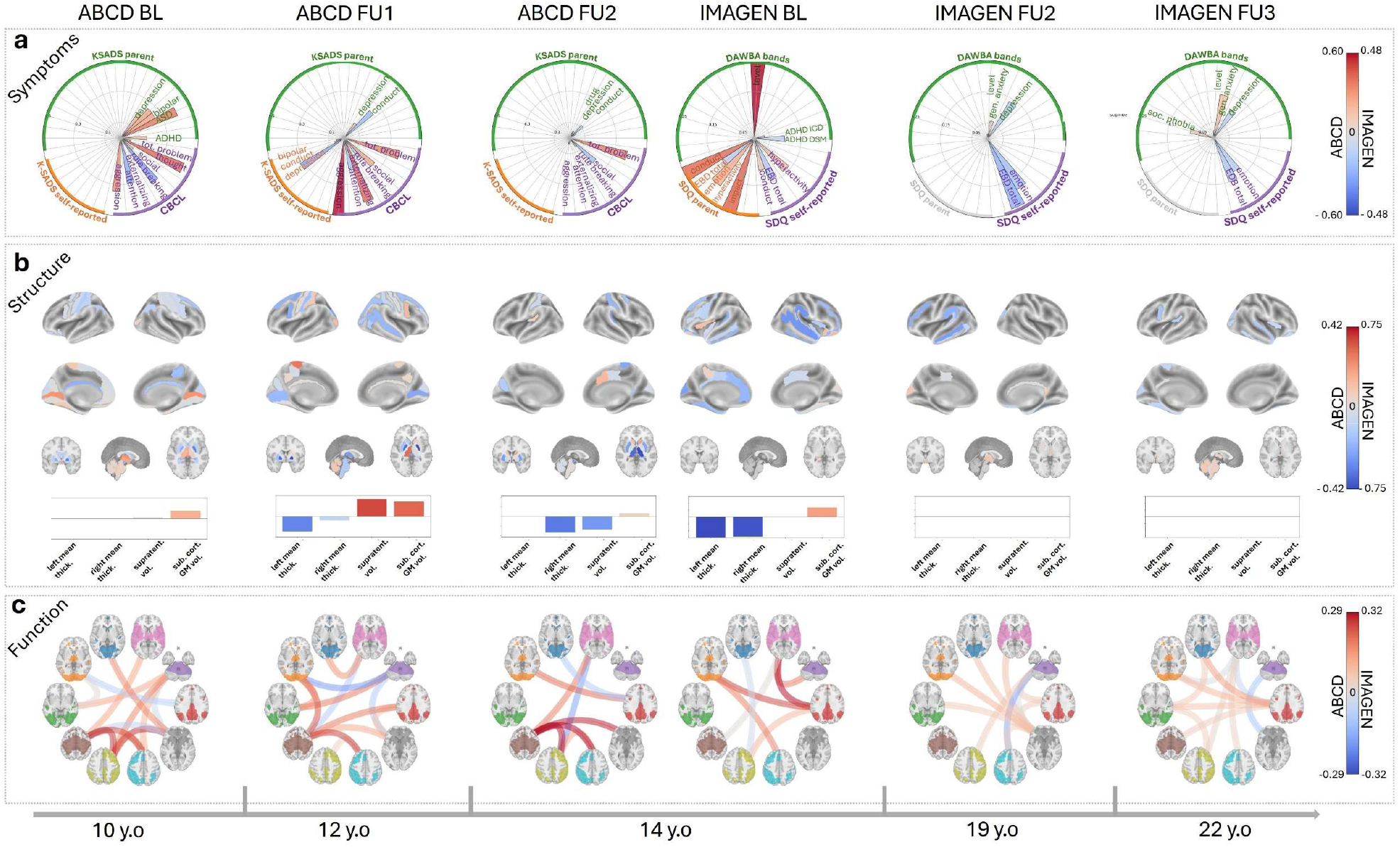
msCCA results over 6 timepoints. **a**, average (over multiple splits) symptom canonical loadings. **b**, average brain-structure loadings. **c**, average loadings in RSN connectivity profiles. for each view, only the stable symptom/IDP scores are displayed (canonical weights that are consistently present 70% of the time across 100 random splits of the data).

### Canonical Loadings and clinical interpretation

To interpret the multivariate brain-symptom relationships identified by multi-view sparse canonical correlation analysis, we present canonical loadings (Pearson correlations between each original variable and its corresponding canonical variate) across our three data views: clinical symptoms, structural neuroimaging measures (cortical thickness and subcortical volume normative z-scores), and resting-state network connectivity z-scores. Canonical loadings provide more direct interpretability than canonical weights: whereas weights serve as coefficients for constructing the canonical variates, loadings directly quantify the strength of association between individual variables and the identified latent dimensions, with the canonical variates themselves being, by definition, maximally positively correlated with one another.

Importantly, our interpretation focusses principally on the patterns of divergence versus convergence between views. Divergence—where loadings from different views show opposite signs—indicates that variables are negatively associated across the multivariate pattern. For example, negative cortical thickness loadings paired with positive symptom loadings would suggest that reduced cortical thickness associates with elevated symptom severity within the identified canonical dimension. Conversely, a convergence where loadings from different views share the same sign indicates that variables covary in the same direction.

We show all features surviving stability selection and their cross-view correlations in supplementary figures S3-5. This can help further understand and confirm the direction and consistency of the results presented next.

### Symptom profiles shift from externalising toward internalising and global measures

Symptoms associated with brain structure showed clear developmental effects. During childhood and early adolescence (ages 10-14), symptom profiles were characterised by prominent externalising features that evolved toward broader psychopathology (Fig. 3a). At age 10, aggressive behaviour and thought problems dominated the positive loadings, while attention problems and rule-breaking loaded negatively, suggesting a mixed externalising profile. At age 12, externalising behaviours remained dominant, with aggressive behaviour being the dominant feature alongside general externalising problems. At age 14, total problem scores became prominent in both cohorts: in ABCD, CBCL total problems and in IMAGEN DAWBA level band in addition to the emotional and behavioural difficulties (EBD) total score from the SDQ, plus other externalising symptoms. This indicates a developmental shift toward global psychopathology rather than domain-specific symptoms.

During mid to late adolescence (ages 19-22), the symptom profile shifts further toward internalising features. More specifically, at age 19 and 22, emotional dysregulation, EBD total scores and generalised anxiety symptoms emerged.

### Structural alterations include widespread cortical thinning and alterations in subcortex

In contrast, the profile of structural alterations associated with transdiagnostic symptom domains remained relatively stable throughout development, although there were some differences at various timepoints (Fig. 3b). The pattern was characterised by a divergent relationship with symptoms in that widespread decreases in cortical thickness predicted an increase in symptoms at most timepoints. Specifically, in ABCD widespread decreases in cortical thickness across lateral temporal, visual and motor regions were predictive of symptoms, accompanied by an increase in overall subcortical volume, largely driven by the cerebellum along with relative decreases in the basal ganglia.

At age 14—the timepoint shared between ABCD and IMAGEN—the structural pattern was again characterised by global decreases in cortical thickness with global increases in subcortical volume, although there were some cohort differences in terms of the specific regions involved. In IMAGEN, the effects were largely cortical, particularly in lateral temporal and cingulate regions. Subcortical effects were only evident globally at 14 in IMAGEN_BL_. Similar global subcortical effects were present in ABCD at the same age. This can be seen as a transition period, where the association was mostly with global symptoms (impact and level-band scores), before transitioning towards internalising symptoms at later IMAGEN timepoints at 19 (IMAGEN_FU2_) and 22 years (IMAGEN_FU3_).

From mid-adolescence onward, alterations became evident in later developing cortical association areas, specifically, the dorsolateral prefrontal (dlPFC) and lateral temporal cortices exhibited negative loadings. In contrast to the early adolescent period (ages 10–14), global cortical thickness no longer contributed stably to the structural canonical variate; instead, a more regionally specific set of cortical structures emerged as correlates of internalising behaviour.

### Functional connectivity transitions from convergence to divergence across adolescence

Resting-state network connectivity demonstrated a complex pattern of associations with symptoms, that provided evidence for developmental effects. Specifically, during early adolescence (10-14), mostly positive network functional connectivity patterns predicted an increase in (externalising) symptoms that later transitioned toward associations between connectivity and internalising symptoms in late adolescence and early adulthood. More specifically, in ABCD, positive connectivity between sensorimotor networks at ages 10-14, the central executive network (age 10) and default mode network (age 14) were predictive of symptoms.

As in ABCD, at age 14 in IMAGEN, default mode network connectivity was also predictive of symptoms although the central executive network was not. At age 19 in IMAGEN, central executive network and default mode network connectivity was predictive of symptoms. At age 22, principally default mode connectivity predicted symptoms.

In order to assist with the interpretation of our findings, we also report the raw correlations between each symptom domain, regional cortical thickness, brain volume and network connectivity (supplementary figures S3–5). This is helpful to facilitate interpretation of our findings, particularly given documented instabilities in CCA models under perturbations to the data or with too few samples per feature^13,28^.

Indeed, the raw correlations between each symptom score and each brain IDP z-score (structural and functional) confirmed the direction of the selected stable features: a positive raw correlation (Fig. S3–5) corresponds to a convergent brain-behaviour relationship, and vice versa.

Next, we performed an additional sensitivity analysis to confirm the directionality of the effects in IMAGEN, which is less straightforward than for ABCD, because not all behavioural instruments were acquired at each timepoint (for instance, the parent-reported SDQ was no longer acquired at ages 19 and 22; Fig S3-4) and because the msCCA model selected different features across each timepoint (Fig 3). Therefore, we performed an additional msCCA restricted to four subject-reported SDQ scores and four DAWBA-bands (i.e. the features that were stable at any timepoint in the main analysis). The results confirm that hyperactivity, social phobia and depression were divergent with brain systems throughout later development (e.g. negatively associated with cortical thickness), whereas generalised anxiety only became divergent in adulthood (i.e. negatively associated with cortical thickness at 19 and 22 years). In contrast, emotion regulation was only divergent in adolescence (i.e. negatively associated with cortical thickness at 14 but not at 19 or 22), which was also evident in the total scores (Fig. S6). Taken together, this: (i) provides further evidence for a shift from externalising/emotional regulation difficulties in adolescence towards internalising difficulties in adulthood and (ii) suggests that sum scores should be interpreted cautiously given that there are many ways that different symptom domains can sum up to give a total score.

Finally, to determine the degree to which our findings are influenced by childhood trauma, we examined whether the stable symptom composite derived from mSCCA was associated with childhood trauma measures. To achieve this, we computed Pearson correlations between all behavioural measures, the stable symptom sum, and trauma scores across cohorts—using the Adverse Childhood Experiences (ACE) proxy score in the ABCD study and the Childhood Trauma Questionnaire (CTQ) in the IMAGEN study (see Methods for details of trauma score derivation). The stable symptom sum was significantly correlated with the overall ACE score across all three ABCD timepoints (r = 0.26, 0.29, and 0.28 at baseline, FU1, and FU2, respectively) and with the CTQ total score across all three IMAGEN timepoints (r = 0.20, 0.28, and 0.26 at baseline, FU2, and FU3, respectively; all p < 0.001).

## DISCUSSION

In this study, we mapped the association between psychopathology, brain structure and function across the entire adolescent developmental period (10-22 years) by aggregating two large population-based cohorts, namely the ABCD and IMAGEN datasets. We report four main findings: first, we found consistent brain-behaviour associations across all timepoints. Second, these associations were largely driven by externalising symptoms during early adolescent timepoints and internalising symptoms during later timepoints. Third, associations with symptoms were driven by relatively consistent and widespread decreases in cortical thickness and more dynamic increases in connectivity. Fourth, the symptom associates we report were strongly associated with childhood trauma. Our results provide evidence for a dynamic and many-to-many mapping between brain organisation and symptoms of mental illness across adolescence.

Our findings broadly align with theoretical and empirical frameworks that posit a shared neural basis underlying psychopathology^6,29,30^, however our results provide a more fine-grained and developmentally grounded perspective that shows how the expression of psychopathology changes over time and is reflected in changes and reorganisation of the core networks. Our results are consistent with numerous studies that also shed light on the brain-behaviour associations using CCA approaches in children^12^, or young adults^15^. The main limitation of these previous studies is that they analysed the brain-behaviour relationship at a specific age or narrow range of the life span. Other studies, investigating brain-behaviour relationship also tend to focus on specific symptoms and behaviours such as adolescents with ADHD or substance abuse symptoms^31,32^. Only few studies have examined the evolution of image-derived brain phenotypes such as cortical thickness or surface area, but did not try to associate those changes to behaviour or symptoms^33,34^. Furthermore, those studies relied on fewer than two hundred scans, with no more than one dataset with maximum three timepoints.

Cortical thickness exhibited predominantly divergent relationships with psychopathology in early adolescence. Lateral temporal, visual, and motor regions showed sustained negative associations from ages 10-14, reflecting structural vulnerability during the transition from childhood externalising behaviours toward global psychopathology^35^. Although there were regional differences across different timepoints, mean cortical thickness remained consistently negative across early adolescence. This global thinning pattern has been widely documented in large-scale studies and likely reflects altered neurodevelopmental processes, including age-related variations in dendritic arborisation and, to a lesser extent, intracortical myelination, in individuals^2,36,37^. After age 14, the dorsolateral prefrontal and temporal cortices emerged as specific structural correlates of emotion-driven symptom scores, whereby thinner cortex in these regions converged with lower emotional and depression scores. This is consistent with evidence that normative cortical thinning in these regions indexes maturation of emotion regulation circuitry: Vijayakumar et al.^38^ showed that thinning of the left dorsolateral and ventrolateral PFC from ages 12 to 16 predicted better cognitive reappraisal at age 19, suggesting that this process reflects refinement of top-down regulatory capacity over anxiety and emotion. Ducharme et al.^39^ further demonstrated that the direction of the cortical thickness–anxious/depressive symptom association reverses across development, with thicker prefrontal cortex becoming associated with more symptoms from age 15 onward. However, this relationship is not uniform across the cortex. Romer et al.^40^ found that thinner temporal pole and left insula in 14–17-year-olds prospectively predicted increases in internalising symptoms, suggesting that while dlPFC thinning may reflect adaptive maturation, temporal pole thinning may instead constitute a structural vulnerability.

Subcortical structures demonstrated a more nuanced pattern, with different deep brain regions dominating at specific developmental stages. During ages 10-14, subcortical patterns predictive of externalising symptoms were characterised by overall increased volume, driven by the cerebellum, but also characterised by decreased basal ganglia volume alongside peak externalising behaviours. This early pattern aligns with evidence that these structures undergo substantial volumetric changes during childhood and are critically implicated in externalising psychopathology ^41–43^. A critical transition occurred at age 14, marked by the emergence of global subcortical grey matter volume as the dominant feature across both datasets alongside thalamic structures showing extreme negative loadings. This convergence is consistent with mounting evidence that mid-adolescence represents a sensitive period for thalamocortical circuit maturation, during which the thalamus plays essential roles in prefrontal cortex development and cognitive function^44,45^. From ages 19-22, the pattern shifted from global subcortical measures toward specific regional involvement. By age 22, the cerebellum re-emerged as being negatively predictive of internalising symptoms, reflecting protracted development of cognitive-affective circuits given the cerebellum’s increasingly recognised roles in emotional regulation, social cognition, and mood disorders^46,47^. The timing of the age 14 transition overlaps with pubertal maturation and reflects fundamental changes in brain organisation during this developmental stage, consistent with evidence that puberty exerts organising effects on subcortical development beyond chronological age^48,49^ and that adolescence represents a period of dramatic subcortical reorganisation creating vulnerability for diverse forms of psychopathology^2^.

Resting-state network connectivity exhibited a more nuanced and dynamic relationship with symptoms relative to structural patterns. During early-to-mid adolescence (ages 10-14), hyperconnectivity between sensorimotor and default mode networks predicted externalising behaviours. This pattern is consistent with observations that immature network segregation and excessive connectivity characterise childhood psychopathology^50,51^. This persistent hyperconnectivity through age 14 suggests incomplete functional network maturation represents a vulnerability factor for transdiagnostic psychopathology. From age 19 onward, connectivity across central executive and default mode networks predicted internalising symptoms negatively, suggesting a developmental divergence from the earlier pattern. This coincides with a period of increasing network segregation, strengthening of long-range connections, and protracted maturation of cognitive control, default mode, and and salience networks^52,53^.

Taken together, our findings suggest that different cortical systems mediate psychopathology risk at distinct developmental windows, with a critical inflection point at age 14 characterised by a switch from externalising to internalising behaviours. This timing aligns with evidence that mid-adolescence represents a sensitive period for thalamocortical circuit maturation^44,45^ and coincides with pubertal transitions that fundamentally alter brain organisation^48^. At the cortical level, this transition is mirrored by increasing regional specificity in thickness–symptom associations. In younger cohorts, cortical thickness accounts for negligible variance in internalising symptoms^54^ and the regions distinguishing internalising from externalising disorders are broadly distributed rather than concentrated in prefrontal or temporal areas^35^. After age 14, this diffuse pattern gives way to regionally specific dlPFC and temporal cortex contributions to the structural profile, paralleling evidence that dissociable dimensions of internalising psychopathology map onto distinct cortical regions in adolescence^55^. In parallel, evidence for developmental transition at age 14 included the shift from specific symptom domains (aggressive behaviour, externalising problems) toward global psychopathology measures (impact and level-band scores) reflecting the increasing comorbidity characteristic of adolescent psychopathology^56^, and intensification of sensorimotor-frontoparietal functional connectivity before subsequent transition to divergence. The convergence of structural, subcortical, and functional transitions at this age parallels the well-documented peak in psychiatric disorder emergence during mid-adolescence^2,57^. Collectively, these lines of evidence identify age 14 as reflecting a pivotal developmental period where brain-symptom relationships undergo fundamental reorganisation across structural and functional domains, suggesting this age as optimal for targeted prevention and early intervention strategies^2^

Finally, the consistent association between the stable symptom sum and childhood trauma measures, across both cohorts and all timepoints, suggests that mSCCA-derived symptom profiles could have meaningful clinical utility. The robust correlations observed (r = 0.20–0.29, all p < 0.001) indicate that early adversity is associated with the most reproducible dimensions of psychopathology at each developmental stage, even as symptom profiles shift across adolescence into young adulthood. Interestingly, adversity-related deviations from normative brain structure in adults have been shown to predict future anxiety^58^, consistent with the internalising-dominant profile emerging in later adolescence in our data. Whether such alterations mediate the trauma-symptom associations observed here across early adolescence to young adulthood remains an open question for future work.

### Study limitations

Several limitations should be acknowledged. First, although we observe developmental changes in both symptom profile patterns and brain structural and functional patterns—and can identify associations between specific symptom profiles and brain profiles at particular timepoints or within defined age ranges— it would be premature to infer that changes in symptom profiles between timepoints are causally linked to concurrent changes in brain structure. Establishing such causal relationships would require analytical approaches that explicitly model within-subject trajectories of change.

Second, the interpretation of RSN connectivity results at age 14 should be approached with caution. In both the ABCD and IMAGEN cohorts, a substantial proportion of resting-state fMRI scans at this age were missing. This missingness poses a challenge for a three-view multi-set canonical correlation analysis (msCCA), as it reduces statistical power, increases the potential for sampling bias, and may lead to unstable canonical variates. Non-random attrition—an inherent issue in longitudinal studies—further compounds the problem, and in ABCD, the COVID-19 pandemic substantially exacerbated data loss and irregularities in acquisition.

Third, behavioural data were not fully harmonised across cohorts. This was a deliberate choice given that: (i) we wished to avoid the well-known difficulties in harmonising instruments, particularly at the item level, and we wished to avoid the dimensionality reduction inherent to most harmonisation methods^27^; (ii) we wished to analyse each timepoint separately. Nevertheless, it is possible that differences in behavioural assessment instruments, scoring methods, and data availability may introduce variability that may affect the comparability and interpretability of results. In particular, our findings suggest that total scores (e.g. the DAWBA level band and the EDB total score from the SDQ) should be treated with caution due to an inherent ambiguity in that many different combinations of symptoms can give rise to the same total score.

Relatedly, our data show that there are residual cohort differences between IMAGEN and ABCD which are a well-documented feature of large cohort studies^59^. Specifically, whilst the overall pattern of effects at the shared 14-year-old timepoint is consistent (e.g. global decreases in cortical thickness predicting a mixed externalising/global symptom profile), there are regional differences underlying these associations between cohorts. This should be taken into account in the interpretation of our findings. To tackle these issues, we are exploring methodological approaches for cross-cohort harmonisation,^27^ and are extending our findings to later ABCD timepoints which are becoming available and to additional cohorts. These results will be reported in future work.

## CONCLUSION

This study employed multivariate canonical correlation analysis across two large, independent cohorts (over 10,000 participants) spanning ages 10 to 22 years to characterise the developmental architecture of brain-behaviour associations in adolescence. Four fundamental patterns emerged, revealing systematic developmental reorganisation rather than static neural correlates of psychopathology.

Symptom profiles evolved from externalising features (ages 10-12) through global psychopathology (age 14) to internalising features (ages 19-22). Cortical thickness exhibited negative associations with externalising symptom profiles during early adolescence (ages 10–14), before shifting to positive associations with internalising symptoms from mid-adolescence onward, accompanied by a transition from diffuse, broadly distributed patterns toward regionally specific involvement of the dorsolateral prefrontal and temporal cortices. Subcortical contributions demonstrated clear temporal specificity: early cerebellum-basal ganglia patterns transitioned through critical age-14 thalamic and global grey matter dominance toward late-adolescent regional contributions. Functional networks transitioned from convergence (cognitive control-sensorimotor hyperconnectivity co-occurring with symptoms) to divergence (network patterns opposing internalising symptomatology), suggesting normative functional maturation despite persistent structural alterations.

These findings challenge the prevailing paradigm seeking static neurobiological markers of psychiatric risk, demonstrating instead that especially functional brain-psychopathology associations undergo systematic developmental reorganisation. The contrast between stable structural divergence and dynamic functional transitions reveals that structural and functional measures capture distinct aspects of neurodevelopmental risk, with functional networks changing through development while structural vulnerabilities persist—providing a substrate for potential symptom remission. This establishes a framework for precision psychiatry recognising developmental stage as a critical moderator of vulnerability, with age 14 emerging as optimal for targeted prevention given convergent thalamic reorganisation, global subcortical dominance, and symptom profile transitions.

## METHODS

### ABCD and IMAGEN datasets

This study used two large and independent datasets: brain imaging and behavioural data from the Adolescent Brain Cognitive Development (ABCD) study^21^ and the IMAGEN study^22^.

In ABCD, we utilised structural T1-weighted MRI scans (sMRI), resting-state functional MRI (rs-fMRI), and the Kiddie Schedule for Affective Disorders and Schizophrenia (K-SADS)^23^ and Child Behavioural Cheklist (CBCL)^24^ measures from around eight thousand young participants over three waves: at 10, 12, and 14 years old (i.e. all data from the ABCD 5.1 release). The K-SADS is a semi-structured diagnostic interview administered to both the child and their parent or caregiver. The K-SADS systematically probes for the presence, severity, and duration of symptoms associated with a wide range of psychiatric disorders in children and adolescents, including mood disorders, anxiety disorders, psychotic disorders, and behavioural disorders, based on DSM criteria. It is designed to capture both lifetime and current episodes, combining clinician judgment with standardised questioning to ensure diagnostic consistency.

In IMAGEN, similar MRI data were used together with the Development And Well-Being Assessment (DAWBA) band scores^25^ (the probabilistic summary outputs from the DAWBA) and the Strengths and Difficulties Questionnaire (SDQ) scores^26^. DAWBA is a hybrid diagnostic tool that combines structured questionnaires, rating scales, and open-ended questions administered to parents, young people, and teachers. Responses are processed using a computer algorithm that generates DAWBA bands—ordinal categories reflecting the probability that a participant meets criteria for specific psychiatric disorders, ranging from very low probability to high probability. This approach allows for both quantitative screening and clinician-reviewed diagnostic confirmation. The SDQ is a brief behavioural screening instrument that assesses five domains: emotional symptoms, conduct problems, hyperactivity/inattention, peer relationship problems, and prosocial behaviour. The SDQ provides both subscale scores and a total difficulties score, offering a concise profile of a participant’s behavioural and emotional functioning. Together, these instruments capture detailed and comparable psychiatric and behavioural phenotypes in each cohort, albeit through different methodologies. There are parent- and self-reported versions of the SDQ. These behavioural/symptom measures were collected from around two thousand participants over three waves: ages 14, 19, and 22. Note that from age 19 onward, SDQ measurements were self-reported only.

Data acquisition was approved by the ethical review boards responsible for the contributing studies: the IMAGEN study received ethical approval from the local ethics committees of all participating sites, and the ABCD study from the central Institutional Review Board at the University of California San Diego. Written informed consent was obtained from all participants and their parents or guardians.

### MRI preprocessing and IDPs

The MRI data were processed using FreeSurfer to allow the extraction of imaged-derived phenotypes (IDPs). In ABCD, we used the processing released by the ABCD consortium, as described in Hagler et al., 2019^60^. Briefly, the T1-weighted images were acquired from 21 sites in the USA (with different scanner vendors), went through the DAIC quality control using a combination of automated and manual methods. Then, the field correction and the resampling to 1mm isotropic voxel size of the data was followed by the freesurfer ‘recon-all’ pipeline, resulting in 150 cortical thicknesses and 31 subcortical volumes. Next, an additional Quality Control (QC) step was performed by experts. For the rs-fMRI, we used an in-house FSL-based pipeline. Briefly, the T2*-weighted (fMRI) images, were registered to their T1-weighted scans. Then, motion correction and resampling to 2.4 mm isotropic voxels in ‘native’ space was applied, followed by automated QC. Next, we denoised these images using ICA-AROMA (non-aggressive denoising). We excluded scans having excessive relative head-movement (> 3mm in any direction), and where the number of ‘good’ networks from ICA-AROMA was too high or too little (thresholded at ± 2.5 SD of the standardised distribution). Finally, resting-state connectomes (pairwise connectivity between the 10 networks from Smith et al^61^) were then generated from these processed data using Nilearn and denoising regression: head-motion parameters (rotations and translations from MCFLIFT; FSL), and physiological noise CSF and White Matter mean signal) and their first derivative as regressors.

In IMAGEN, data were acquired from 8 sites in Europe (France, UK, and Germany), and similar preprocessing steps were taken. Rs-fMRI data were processed using an fMRI-prep pipeline^22^, which includes FreeSurfer. Data were also realigned and subjected to ICA-AROMA (non-aggressive denoising) and normalised to a standard template using the ANTS routines. Then, additional QC was applied following the steps taken in ABCD, (visual inspection, > 3 mm head-movement, and IC networks outliers) and calculated the resting-state connectomes using Nilearn and denoising regressions (same regressor as in ABCD connectome calculation). For the sMRI, the preprocessed T1w from FreeSurfer were used, and manual QC (visual inspection of fMRI-prep reports) performed. 150 cortical thickness and 38 subcortical volumes IDPs were extracted. To align the data to ABCD, six subcortical volume IDPs were removes from the analysis, namely the estimated total intracranial, left and right choroid plexus, left and right vessel, non-ventricular supratentorial, and total grey matter volumes.

All cortical thicknesses, subcortical volumes, and all pairwise connectivity measure from resting-state connectomes represent the structural and the connectivity IDPs. These were then fed to the normative models in order to extract the normative deviation z-scores, as presented in the next section.

### Normative modelling and image-derived phenotypes

Structural measurement and functional data from MRI are known to be highly dependent on (the non-linear effect of) participants’ age and the acquisition site. The best way to controls for those covariates, and to obtain robust imaging-derived phenotypes (IDPs) is the use of normative models^17,19,62^ (NMs).

The baseline (BL) ABCD resting-state and structural MRI data were already used in the pre-trained NMs^18^ Therefore, we just predicted the NM deviation scores (z-scores) directly for BL (participant aged 10), follow-up 1 (FU1; at age 12), and follow-up 2 (FU2; at age 14) timepoints. More specifically, we used the Cortical Thickness and Subcortical Volume Bayesian Linear Regression (BLR) model (58,836 samples; 82 sites) and the fMRI Smith-10 Resting-State Brain Network BLR model (21,515 samples; 45 sites) for the NM deviation score predictions of the structural and functional IDPs, respectively.

In the IMAGEN dataset, the predictions of z-scores were done using the transfer function of the PCNtoolkit (https://www.github.com/amarquand/PCNtoolkit) which utilises a small sample to first augment the NMs in order to model the effects of new IMAGEN sites and age. Specifically, we used 20 participants from each of the 8 sites as adaptation data in both NMs. Predictions are then made on the rest of the data, using the same pre-trained models mentioned above. Only these newly predicted NM z-score data are used for further canonical correlation analysis (CCA). This is performed on all three waves of IMAGEN data collection: BL (age 14), FU2 (age 19), and FU3 (age 22). Note that FU1 (age 16) was skipped since it contains only behavioural and cognitive assessment measurements.

The resulting sample sizes in ABCD, after preprocessing and NM were: baseline (BL, age 10) 7,961 / 7,961 / 7,961 for behavioural, structural, and resting-state connectivity data, respectively; first follow-up (FU1, age 12) 5,373 / 5,373 / 5,373; and second follow-up (FU2, age 14) 1,806 / 1,806 / 672. For the IMAGEN cohort, the corresponding sample sizes were: BL (age 14) 1,860 / 1,860 / 265; FU2 (age 19) 985 / 985 / 824; and FU3 (age 22) 983 / 983 / 809.

### Analysis pipeline

Figure 1 depicts the analysis pipeline for our multi-view sparse canonical correlation analysis (msCCA). Data were first preprocessed and QCed as described in the previous sections (Fig. 1A). A randomised split of the processed data (IDP z-scores and behavioural scores) was applied to select the training (70%) and testing (30%) samples. We then trained the multi-view sparse CCA using the IDP z-scores and behavioural scores (see next section). Using the default 0.5 sparsity/penalty parameters for each of the views, as in prior work^12,15^, the trained msCCA models were then fit to the unseen (testing) data. This analysis was repeated 10 times with 10 different splits of the data for ABCD data and 30 times for the IMAGEN dataset. After averaging the resulting canonical correlations between the brain and behavioural metrics, significance testing was performed on both the in-sample (training) results and the out-of-sample (validation) averaged correlation using permutation testing. The higher number of splits for IMAGEN was chosen to account for the greater variance across random draws in smaller samples, where more repetitions are needed to ensure a stable and representative averages.

In parallel, we ran stability selection to identify consistent IDP and behavioural weights. This involved repeating the msCCA 100 times, using random splits of the data and identifying weights that were selected in more than 70% of the splits. This threshold is motivated by stability selection theory^63^ as it provides guarantees over the family-wise error rate. Finally, for each view of the data (cortical thickness or volume, network connectivity and symptom scores) these stable brain and behavioural phenotypes were extracted and the canonical loadings were computed (Fig. 3).

It is worth noting that the sparsity parameter of 0.5 was selected after a thorough MSCCA optimisation search performed on both the ABCD and IMAGEN datasets, as in Mihalik et al., 2020^64^. In both cohorts, the optimised sparsity parameters did not lead to significantly better canonical correlations in the testing sets than the default of l1 = 0.5. More information on the optimisation analysis can be found in the Supplementary Information.

### Multi-view Sparse Canonical Correlation Analysis

After predicting NM z-scores for the MRI data in ABCD and IMAGEN, we assessed the following correlations: Behaviour scores with structural MRI z-scores; Behavioural scores with rs-fMRI z-scores; and Structural MRI z-scores with rs-fMRI z-scores. We applied the Multiview Sparse Canonical Correlation Analysis (MSCCA) as in implement in Ing et al.^15^ In short, this optimises for maximal correlation between view1-view2, and view1-view3, by maximising the following relationship:

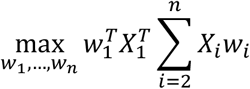

Subject to the constraints:

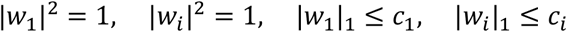

This method can be viewed as a hierarchical or semi-partial CCA, in that the covariance between view2-view3 is ignored (which is crucial to avoid learning trivial correlations between imaging modalities), and view1 is viewed as the hub, and the emphasis of the analysis is on finding the shared modes variation between view 1 (symptoms) and the other brain dimensions (brain structure and function). Since it is known that brain structures and functions are highly correlated, optimising the CCA weights for that correlation would bias the resulting predictions, and most likely underfit them.

### Statistics and significance testing

To test the reliability of the canonical correlations, we performed 10 randomised splits of the data (70% training, 30% testing) and averaged the results. However, since we cannot assume full normality in the distribution of the correlation values, non-parametric permutation testing was used to assess significance. Specifically, we permuted the first view (symptoms) of the data, performed the 10-split (or 30-split) analysis, and repeated this 100 times. This yielded a 1,000-/3000-permutation null distribution of two pairs of canonical correlations (view1 with view2 and view1 with view3). We then averaged the correlation between these two pairs, and effectively obtained the overall brain-behaviour canonical correlations null distribution. By comparing our averaged canonical correlations from the 10 splits against this null distribution, we obtained p-values for the MSCCA results. The significant canonical correlation (at p < 0.05) of each timepoint are depicted in Fig. 2.

### Childhood trauma measures

In IMAGEN, childhood trauma was assessed using the short form of the Childhood Trauma Questionnaire (CTQ)^65^, administered at FU2 (age 19) and applied to all three IMAGEN timepoints. CTQ subscale scores (emotional neglect, emotional abuse, sexual abuse, physical abuse, physical neglect) and the CTQ total score were used. In ABCD, childhood adversity was quantified using a youth-deferred ACE proxy score derived from the ABCD 5.1 release following Breslin et al.^66^, using youth report only. Two summary scores were computed: a 5-domain CTQ-matched proxy (0 to 5) restricted to physical abuse, sexual abuse, emotional abuse, physical neglect, and emotional neglect, corresponding directly to the five CTQ subscales to enable cross-cohort comparability; and a full 10-domain ACE score (0 to 10) additionally encompassing household dysfunction domains (parental separation, domestic violence, mental illness in the family, incarceration of a family member, and family substance use). Both scores were accumulated prospectively across waves (baseline through year 4), with earlier endorsements carried forward to subsequent timepoints. Pearson correlations were computed between all behavioural measures, the trauma scores, and the stable symptom sum—defined as the unweighted sum of stable symptoms identified through the mSCCA stability analysis at each timepoint—to assess the relationship between trauma history and the most reproducible features of the symptom profiles.

## Author contributions

Resources & Data curation: T.B., A.L.W.B., R.B., S.D., H.F., H.G., P.G., A.G., A.H., H.L., J.L.M., M.L.P.M., E.A., F.N., D.P.O., T.P., L.P., M.N.S., N.H., N.V., H.W., R.W., P.W., G.S

Conceptualization & Methodology: A.B., A.M.

Investigation & Visualization: A.B.

Supervision: A.M.

Writing—original draft: A.B., A.M.

Writing—review & editing: A.B., A.M., L.S., T.B, T.P, A.H.

## Data, code, and materials availability

Data used in the preparation of this article were obtained from the Adolescent Brain Cognitive Development^sm^(ABCD) Study (https://abcdstudy.org), held in the NIMH Data Archive (NDA); and from the IMAGEN study (https://www.imagen-project.org/). Please visit the respective websites for information on how to access the data. All code used to perform the analyses in this paper is available at: https://github.com/predictive-clinical-neuroscience/SCCA_adolescence

