## Supplementary information for "Trajectories of brain organisation transition from predicting externalising to internalising symptoms across adolescence"

Antoine Bernas *et al.*

**This PDF file includes:**

Supplementary Text
Figs. S1 to S6

**Supplementary Text**

msCCA second mode

Our main results in the manuscript focused on the first, and most important, model order of the multi-view sparse canonical correlation analysis (mSCCA), i.e., the first mode. However, the second mSCCA mode showed statistically significant correlations between the views at all time points/waves (Fig. S1).

Even though the statistics were significant, only a few canonical weights were stable (more details in Materials and Methods). This is evident when examining the symptoms and RSN functional connectivity highlighted in Fig. S2a. For example, in the third wave IMAGEN FU3, symptom scores and functional IDPs had no stable canonical weights. Additionally, when weights were stable, their structure coefficients (or loadings) were in the low range, reaching a maximal Pearson correlation value of 0.27 in ABCD BL time point for structural features, as compared to 0.75 in some cases in the first CCA mode (see Fig. 3, main text). Therefore, the results depicted in Fig. S2 must be interpreted with caution.

Nevertheless, several remarkable properties of the mSCCA method and brain-behavior associative evolution in adolescents can be observed: (i) the second mode highlights complementary variables (those not typically prominent in the first mode), revealing secondary brain-behavior associations that occur as less dominant effects; for example, in ABCD FU2 (14 y.o.), self-reported K-SADS anxiety-related scores correlated positively with lateral and inferolateral ventricle volumes. (ii) The second mode can also highlight the same behavioral, structural, or functional variables as in mode 1, but the association may be reversed—that is, diverging when the association was converging in the first canonical mode. For instance, in the ABCD FU1 second canonical mode, left mean cortical thickness now converges with bipolar and depression scores, indicating that thinner overall left cortex is associated with reduced internalizing symptom severity. While the second observation (ii) may appear to contradict findings in the first mode, it often underscores that results should be assessed within a more global framework: symptom "profiles" are associated with brain "profiles," not individual variables, emphasizing the necessity of multivariate analysis. Additionally, observation (ii) can place even greater importance on primary findings: indeed, in the brain-behavior associations in ABCD FU1, the primary findings demonstrate that, at that age, mean cortical thickness and mean subcortical volumes are associated with externalizing symptoms (see Fig. 3, main text), while internalizing symptom associations are more complex and secondary.

Finally, what we consider the most interesting finding in the second mode of CCA models is the combination of (i) and (ii) at different time points—that is, the same variable in one view showing an opposite (relative to the first mode) association with complementary variables/profiles of another view at a different stage of adolescence. For example, in IMAGEN FU2 mode 2, strong negative loadings of mean cortical thickness reemerged (previously the strongest loadings in IMAGEN BL mode 1), but this time they were associated with a simplified symptom profile comprised of only one stable symptom, the SDQ Prosocial score, in a convergent manner (positive association). Since the prosocial score represents a positive value (higher scores indicate better mental health) and is essentially complementary and opposite to the SDQ EDB total score, the shift in direction of the association is coherent. Hence, such canonical correlations can provide stronger evidence of brain-behavior associations that are critically important at certain time points and subsequently become secondary later in development.

Raw data (structure, function, and symptoms) correlations

Figs. S3-S5 depict the correlations between raw data at each time point. It is evident that correlations between stable symptoms (in terms of canonical weights—see Methods) and brain IDPs (structural or functional) remain consistent across time points, i.e. predominantly maintaining their sign. This renders our results regarding the direction (positive/convergent or negative/divergent) of brain-behavior associations robust and consistent across our models derived independently at different time points.

Supplementary constrained msCCA analysis

Fig. S6 shows the constrained msCCA in which only symptom scores consistent across timepoints were used, namely the subject-reported SDQ scores of conduct, total emotional and behavioral difficulties (EBD total), emotion, hyperactivity, and social phobia, as well as the DAWBA-bands for total level, general anxiety, and depression. The changes in sign of the EBD total, emotion, and general anxiety scores from age 14 to ages 19/22 corroborate our main finding of a shift in the direction of structure-symptom associations, with the temporal and dlPFC cortices emerging as regionally specific correlates after age 14. Similarly, the constrained msCCA provides converging evidence that the same internalizing symptoms diverge with functional connectivity dominated by the central executive network at age 19 and the default mode network at age 22, consistent with the patterns reported in the main analysis (Fig. 3c)

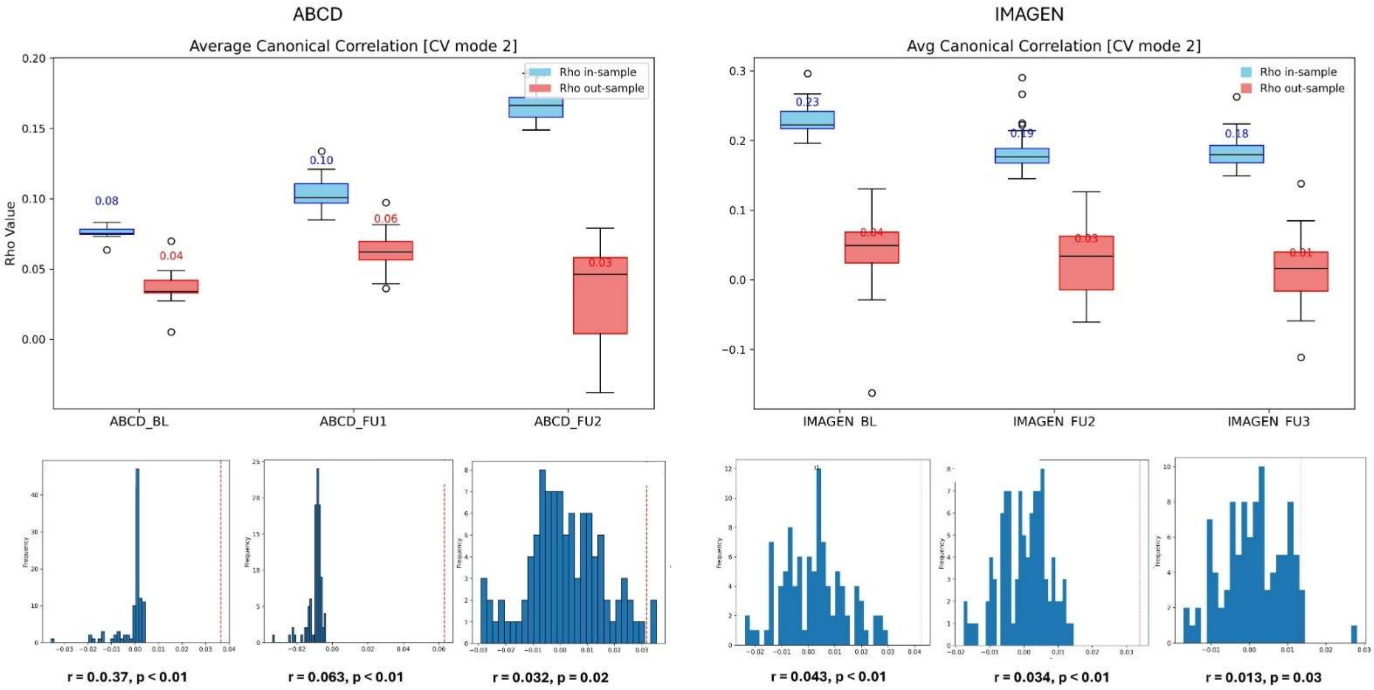


**Fig. S1. Significant canonical correlations, 2nd Mode.** Average (30-split) canonical correlation between the behavioral and brain-related canonical variates (top); and canonical correlation null distribution (bottom); for $ABCD_{BL}$, $ABCD_{FU1}$, and $ABCD_{FU2}$, (left); and $IMAGEN_{BL}$ , $IMAGEN_{FU2}$ , and $IMAGEN_{FU3}$ (right); In-sample (training) results in blue and Out-of-sample (test) results in red. The numbers displayed on the boxes are the 30-split average correlations, while the boxes show the usual 25th, median, and 75th centiles. Hollow circles = outliers, when correlation is outside of the whiskers (interquartile range x 1.5).

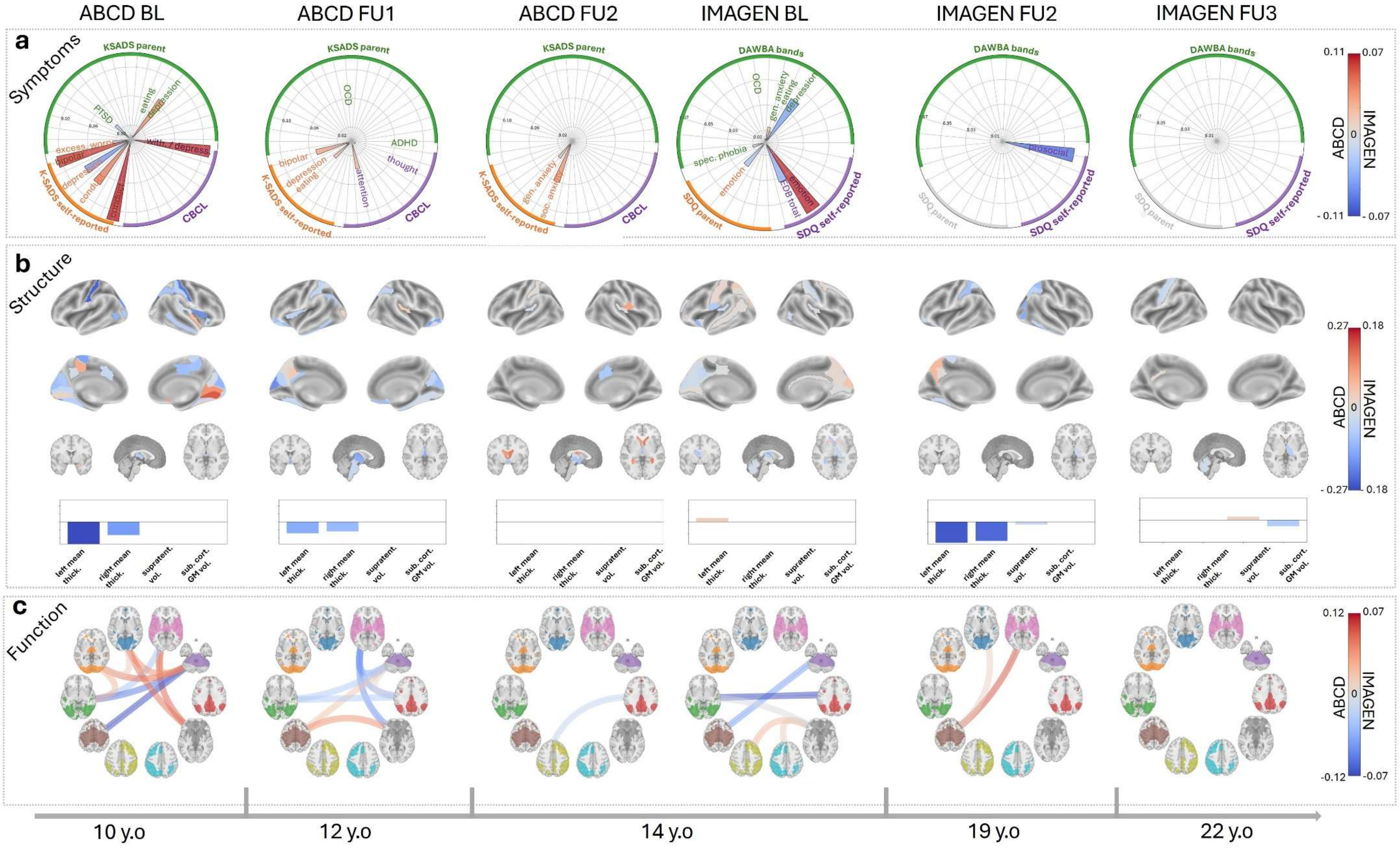


**Fig. S2. MSCCA results over six timepoints, 2nd Mode. a**, average (30-split) symptom canonical loadings. **b**, average brain-structure loadings. **c**, average loadings in RSN connectivity profiles. for each view, only the stable symptom/IDP scores are displayed (canonical weights that are consistently present 70% of the time in 100 MSCCA).

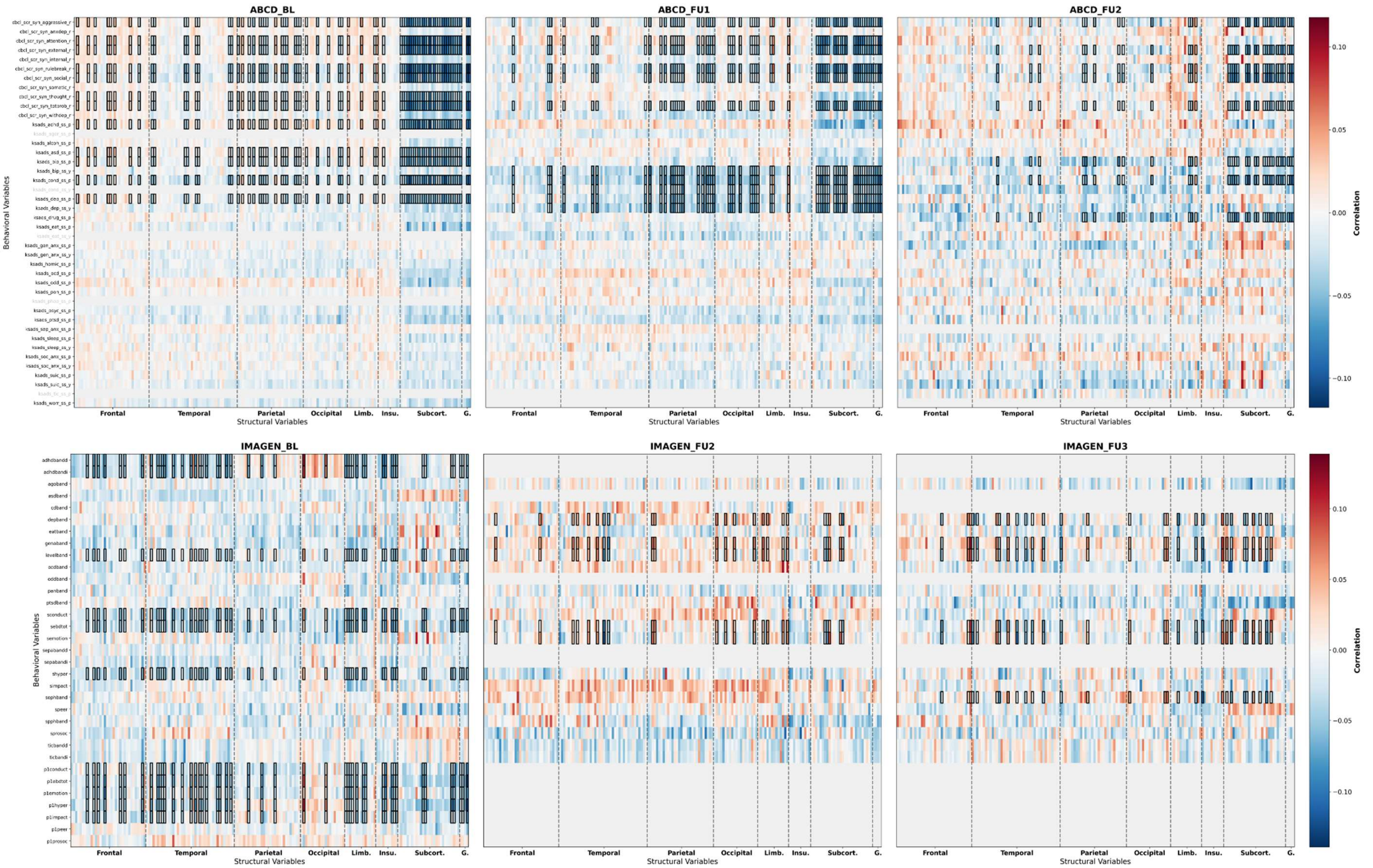


**Fig. S3. Symptoms-to-structure correlation across the six timepoints. a**, ABCD's correlation between raw symptom scores and structural Normative Model (NM) Imaged-derived phenotype (IDP) zscores **b**, IMAGEN's correlation between raw symptom scores and structural NM IDP zscores. Y-axis = behavioral data; X-axis = cortical thicknesses and subcortical volumes; Highlighted boxes = features that had 70% stable MSCCA weights; Grey zones = missing data, i.e. no symptom scores (rows) in some time points.

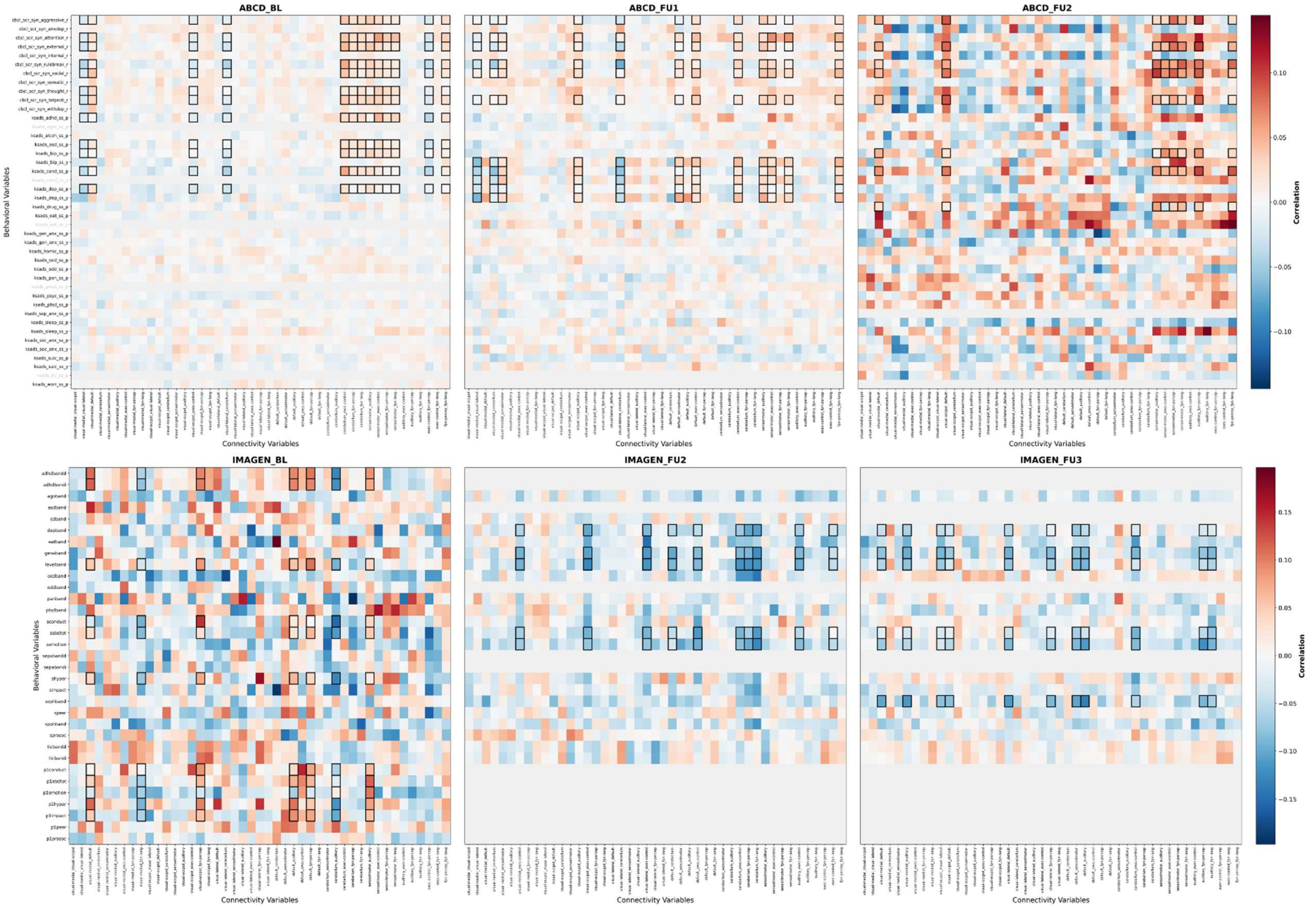


**Fig. S4. Symptoms-to-connectivity correlation across the six timepoints. a**, ABCD's correlation between raw symptom scores and resting-state network (RSN) functional connectivity (fc) NM IDP zscores **b**, IMAGEN's correlation between raw symptom scores and RSN-fc NM IDP zscores. Y-axis = behavioral data; X-axis = RSN pairwise connectivity data; Highlighted boxes = features that had 70% stable MSCCA weights; Grey zones = missing data, i.e. no symptom scores (rows) in some time points.

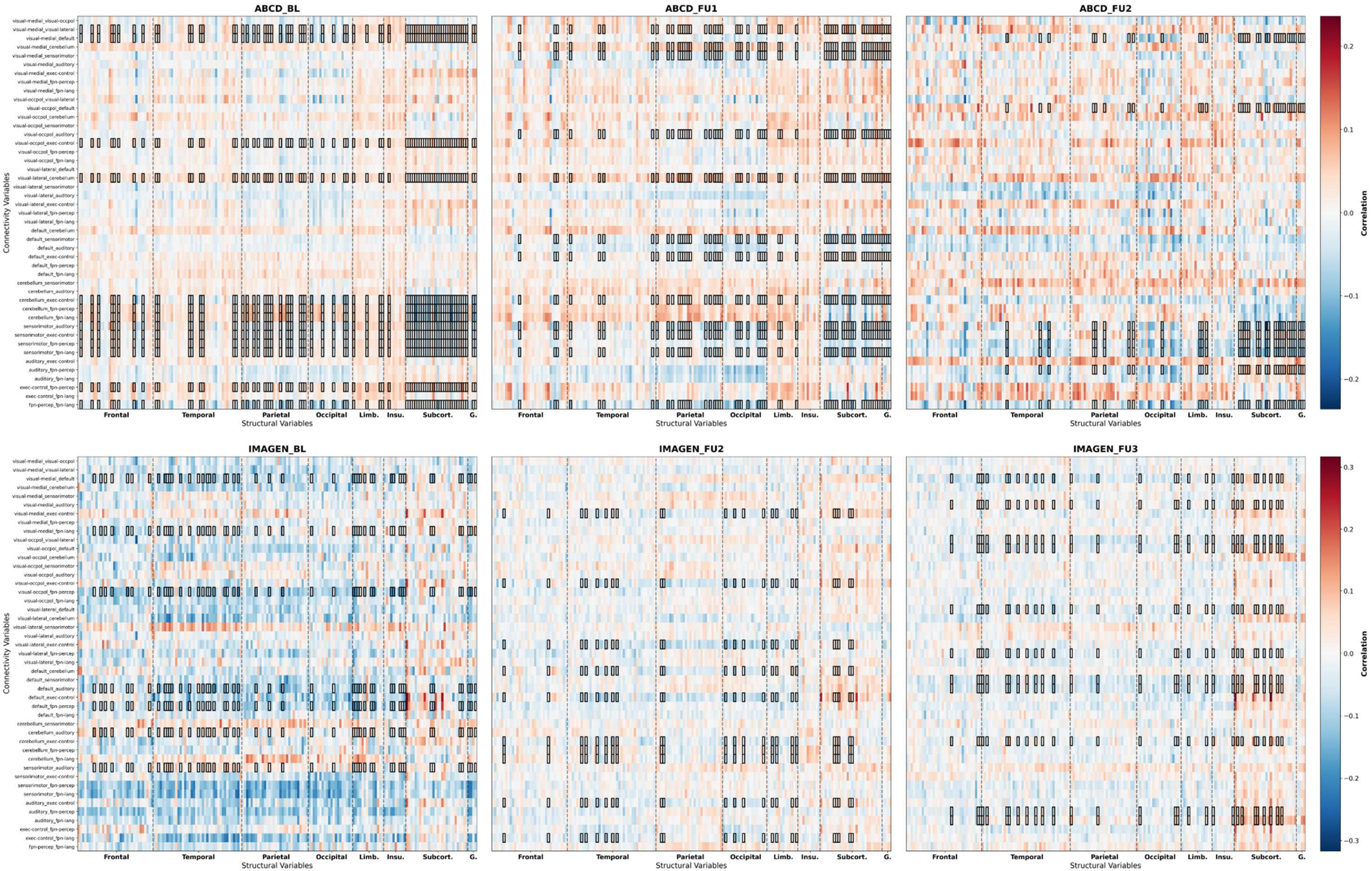


**Fig. S5. Correlation between function and structure across the six timepoints. a**, ABCD's correlation between structural IDP zscores and RSN-fc IDP zscores **b**, IMAGEN's correlation between structural zscores and RSN-fc IDP zscores. Y-axis = RSN pairwise connectivity; X-axis = cortical thickness and subcortical volume data. Highlighted boxes = features that had 70% stable MSCCA weights

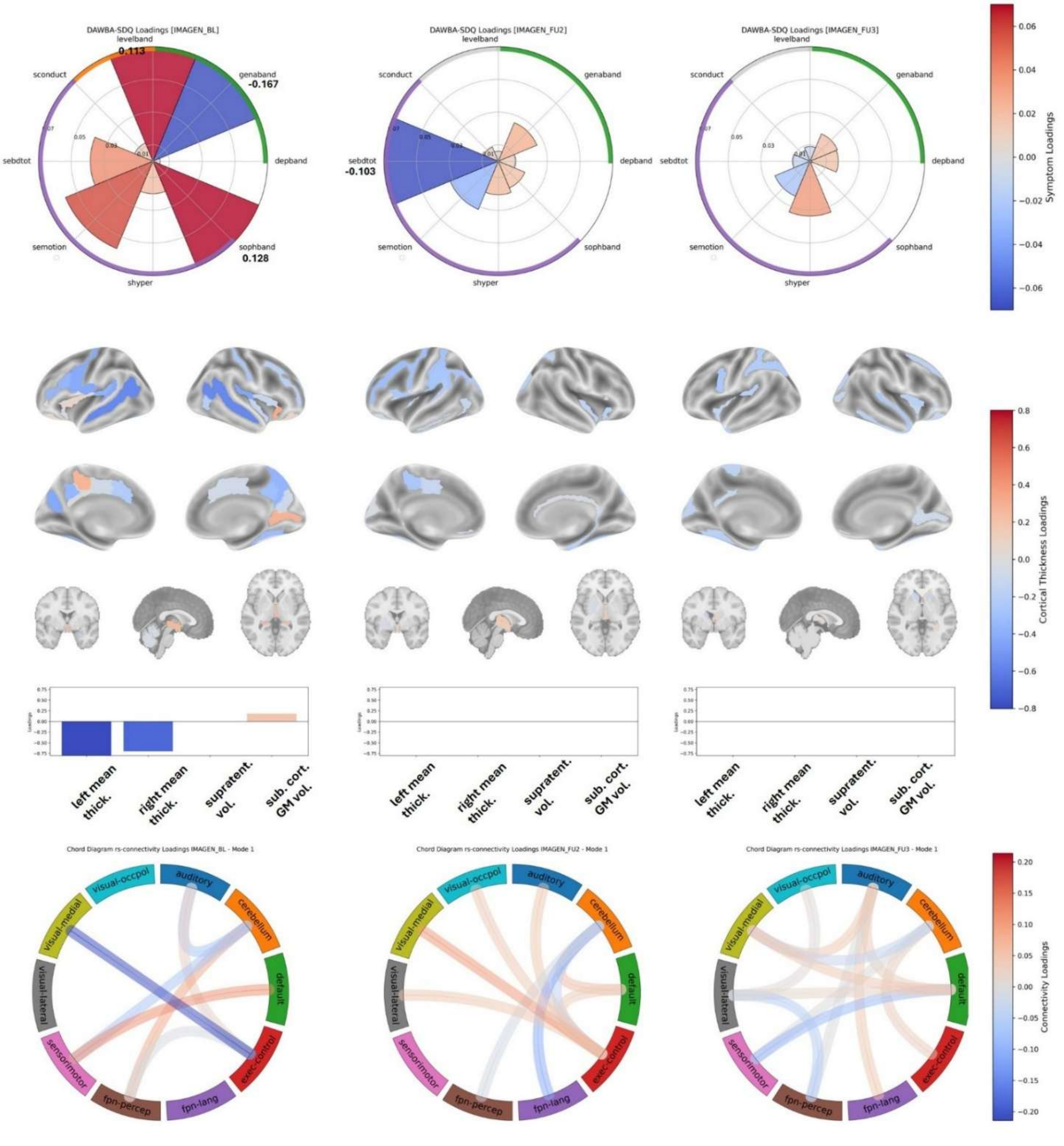


**Fig. S6. Symptom-constrained msCCA in IMAGEN.** MSCCA results over the 3 time points of IMAGEN (age 14, 19, and 22). Average (over 30 splits) of the 8 common symptom canonical loadings (top); Average brain-structure loadings (middle); and average loadings in RSN connectivity profiles (bottom). For the structural and functional views, only the stable symptom/IDP scores are displayed (canonical weights that are consistently present 70% of the time across 100 random splits of the data). *sconduct*, *sebdtotal*, *semotion*, *shyper* are subject-reported conduct, total emotional and behavioral difficulties, emotion, and hyper-activity SDQ scores respectively; and *sophband*, *levelband*, *genaband*, and *depband* are the social phobia, total level, general anxiety, and depression DAWBA-bands respectively.
